# TransXplorer: an automated translational discovery platform for RNA-seq data

**DOI:** 10.64898/2026.05.15.724657

**Authors:** Varinder Madhav Verma, Eponine Oler, Hussain Syed, Scott Han, Mark Berjanskii, Andrew L. Mason, David Wishart, Gane Ka-Shu Wong

## Abstract

RNA-seq experiments routinely identify thousands of differentially expressed genes, but translating these into biological insights and therapeutic hypotheses often requires integrating multiple tools. Existing web platforms such as iDEP, NetworkAnalyst, and GEPIA2 address individual steps, differential expression, network visualization, or TCGA queries, but lack a unified environment spanning raw data processing to clinical and pharmacological interpretation.

TransXplorer (https://transxplorer.org) is a freely available web platform that addresses this limitation by integrating the complete RNA-seq analytical workflow. It supports processing from raw FASTQ files using HISAT2 or Salmon, as well as direct GEO dataset import with automated metadata handling. Differential expression analysis is implemented via DESeq2, edgeR, and limma-voom, followed by functional enrichment across more than 1,800 species using Bioconductor resources. Batch effects are automatically detected and corrected using a composite of PVCA, kBET, and Silhouette metrics without requiring predefined batch annotations. Downstream analyses include co-expression network construction (WGCNA), protein-protein interaction mapping (STRING), cell-type deconvolution, and transcription factor inference using integrated DoRothEA and TFLink resources. The platform further links gene signatures to drug candidates through DGIdb and OpenTargets and enables survival and tumour-normal comparisons across TCGA cohorts.

Application to cardiac endothelial differentiation (GSE151427) and kidney renal papillary cell carcinoma (TCGA-KIRP) datasets demonstrates accurate batch correction, biologically consistent pathway enrichment, recovery of expected cell-type proportions, and identification of clinically relevant genes and drug candidates. TransXplorer is freely available without a login.

## INTRODUCTION

In biomedical research, RNA sequencing has emerged as a widely used technique for transcriptome profiling. The entire pipeline from raw data to quality control, differential expression, functional interpretation, network inference, clinical validation, and pharmacological annotation is currently not covered by a single web-based platform, even though individual analytical phases are well-established. When data needs to be reformatted between platforms, researchers usually assemble different tools for each stage, which deters experimental biologists who do not write code and creates discrepancies (1,2). There are many interrelated analytical gaps, each only partly addressed by existing tools, that illustrate the problem.

Data accessibility is the first. Reanalyzing public RNA-seq from the NCBI Gene Expression Omnibus (GEO) (3). It typically requires manual downloading, format parsing, and metadata setting, and bench scientists often lack access to the command-line alignment and quantification pipelines needed to start from raw FASTQ. Existing web applications such as iDEP(4) and GEPIA2 (5), only accept pre-processed count matrices or focus on pre-computed TCGA data, leaving users without local bioinformatics infrastructure unsupported.

Batch effect confounding is the second. Technical variation introduced during sample processing can obscure or mimic biological signals (6). Most platforms either require users to identify batch variables manually, assuming familiarity with the experimental design, or ignore batch effects entirely; tools like iDEP (4) and DEBrowser (7) offer optional correction but no automatic detection, so non-specialists may miss confounding altogether.

Limited organism coverage is the third. Most web platforms restrict functional enrichment and gene annotation to humans and mice, leaving out comparative genomics, agricultural, and ecological research that increasingly relies on non-traditional model organisms.

Regulatory network inference is the fourth. Identifying the master transcription factors driving differential-expression programs reveals upstream regulators that coordinate large downstream changes and offer mechanistic insight beyond gene lists; gene regulatory networks (GRNs) capture these directed TF → target interactions (8). However, web-accessible curated TF-target databases remain restricted to humans and mice, leaving most model species without this analysis, even where platforms like NetworkAnalyst provide network capabilities (9).

Translational mapping is the most important. Converting gene-level statistical results into drug-target hypotheses requires querying multiple pharmacological databases, cross-referencing clinical trial evidence, and evaluating target druggability, skills outside most non-pharmacoinformatics labs, and this stage is not integrated by any current RNA-seq web platform.

All of these gaps are filled in a single environment by TransXplorer. It allows for the direct import of GEO datasets with automated metadata parsing and an optional FASTQ processing pipeline that uses Salmon and HISAT2. It employs a quantitative scoring approach for batch detection that assesses each metadata variable using three complementary metrics with thresholds that have been empirically calibrated. It incorporates Bioconductor’s AnnotationHub for organism coverage, including functional enrichment for more than 1,800 species. For regulatory analysis, it uses a hybrid inference method that combines projected TFLink relationships (10) across 11 model species with curated DoRothEA transcription factor-target interactions (11). It offers a real-time query engine for translational mapping that creates cross-validated drug-target rankings with clinical evidence by mapping differentially expressed genes against DGIdb (12) and OpenTargets (13). These features offer a comprehensive analytical workflow from raw data to biological and therapeutic interpretation, along with differential expression, pathway enrichment, co-expression network analysis, protein-protein interaction networks, cell-type deconvolution, and TCGA clinical validation across 33 cancer cohorts.

## METHODS

### Input data and FASTQ processing

TransXplorer accepts gene expression count matrices with matching sample metadata (CSV, TXT, TSV, XLSX). Paired-end and single-end reads from seven reference genomes (hg38, mm10, rn6, dm6, danRer11, wbcel235, r64) are supported by an optional FASTQ processing pipeline with two quantification strategies: Salmon (14) pseudo-alignment with tximport (15) for transcript-level quantification aggregated to the gene level, and HISAT2 (16) genome alignment followed by featureCounts (17) for gene-level counting. Compared to the genomic alignment method, salmon quantification is around ten times faster. Quality control includes FastQC (18) reports before and after Trimmomatic (19) adapter removal. Custom genomes are supported through user-provided FASTA and GTF files. Processing runs in the background with real-time progress monitoring, a job queue manages concurrent analyses, and results are downloadable as count matrices or complete result archives.

### Direct GEO dataset import

To simplify access to public transcriptomic data, TransXplorer enables direct import of datasets from the NCBI Gene Expression Omnibus (3). Users enter a GEO Series accession (e.g., GSE164073), and the platform retrieves sample metadata, parses experimental group annotations from GEO’s structured metadata fields, and extracts count matrices through a multi-strategy approach. The platform first attempts to download NCBI-precomputed RNA-seq counts generated through the standardized HISAT2/featureCounts pipeline; if unavailable, it searches supplementary files for count data; as a final fallback, it examines the GEO expression matrix. Extracted metadata is automatically parsed into clean experimental variables (e.g., tissue type, treatment condition) and presented in an interactive configuration panel where users select the primary grouping variable. All remaining metadata columns are retained for automatic batch effect assessment, so no separate metadata file upload is required.

### Differential expression analysis

Three statistical frameworks are available for identifying differentially expressed genes: DESeq2 (1), which models count data using negative binomial distributions with empirical Bayes shrinkage; edgeR (20), employing quasi-likelihood F-tests for robust dispersion estimation; and limma-voom (2), which applies precision-weighted linear models after log-CPM transformation and mean-variance modelling. Multiple normalization methods are supported, including TMM, RLE, VST, CPM, and TPM, each guided by the selected statistical framework.

### Quantitative batch effect detection and correction

TransXplorer evaluates every metadata column in the uploaded sample annotation using three complementary metrics, each measuring a different aspect of batch confounding.

1. **Principal Variance Component Analysis** (PVCA) determines the percentage of overall variation related to each metadata variable by applying a linear mixed-model framework to the expression matrix’s leading principal components. A variable is marked if it contributes more than 15% of the overall variation. This cutoff was calculated using publicly accessible benchmark datasets with known batch structure (GSE49712, GSE115736), where biological covariates of interest stayed below the cutoff but confirmed batch variables regularly exceeded 15%. Additionally, published variance partitioning recommendations for RNA-seq quality control are in line with the 15% cutoff (6).
2. The **k-nearest Batch Entropy Test** (kBET) (21) compares the batch composition of each sample’s immediate neighborhood to the global distribution to determine if samples from various batch categories mix equally in expression space. Rejection rates greater than 0.3 indicate batch-driven segregation.
3. **Silhouette coefficients** quantify the degree to which samples cluster according to batch label rather than biological state. Normalized scores greater than 0.15 suggest that sample clustering is caused by batch identity rather than experimental design.

These measures are combined into a weighted composite (0.4 × PVCA + 0.4 × kBET + 0.2 × Silhouette). PVCA and kBET receive equal weight because they probe complementary aspects, global variance structure and local sample mixing, each of which can miss confounding the other detects. Silhouette is down-weighted because clustering geometry is sensitive to unbalanced group sizes. The composite threshold of 0.25 was tuned so that variables exceeding it on at least two of three metrics are flagged; on calibration datasets, this correctly identified known batch variables without false positives from biological covariates. When a variable exceeds the threshold, limma’s removeBatchEffect (2) is automatically applied, and the variable is included as a covariate in subsequent differential-expression models. PCA and UMAP projections are generated before and after correction for visual confirmation.

## Functional analysis

### Pathway enrichment and protein interaction networks

Gene Ontology (biological process, molecular function, cellular component), KEGG, Reactome (22), WikiPathways, and MSigDB Hallmark gene sets are all supported by functional enrichment using clusterProfiler (23). The entire collection of tested genes is employed as a statistical background for both gene set enrichment analysis (GSEA) and over-representation analysis (ORA). Functional links between differentially expressed gene products are shown using STRING-based (24) protein-protein interaction networks, which leverage locally stored data for human and mouse (removing API rate limitations). Hub proteins and functional modules are identified by Louvain clustering using network topology criteria, including degree, betweenness, proximity, and eigenvector centrality. The platform additionally ranks hubs using the 11 CytoHubba (25) algorithms (MCC, MNC, DMNC, EPC, BottleNeck, EcCentricity, Closeness, Radiality, Betweenness, Stress, and Degree), enabling consensus hub identification across complementary topology criteria.

### Auto-optimized co-expression network analysis

Weighted gene co-expression network analysis (WGCNA) (26) requires selecting a soft-threshold power parameter to determine the network topology. By utilizing the pickSoftThreshold function to evaluate powers ranging from 1 to 20, TransXplorer automates this process and chooses the lowest power that achieves a scale-free topology fit (R^2 >= 0.8). Hierarchical clustering with dynamic tree cutting is used in module discovery. The first main component of each module’s expression matrix, known as module eigengenes, relates to sample characteristics and summarizes overall expression profiles. The platform performs functional enrichment on each module to assign biological categories and computes module membership (kME) and gene significance scores to discover hub genes.

### Multi-algorithm cell-type deconvolution

Three deconvolution methods estimate cell-type proportions in mixed-tissue samples, each suited to a distinct biological setting: MCP-counter (27) estimates the absolute abundance of ten immune and two stromal populations; EPIC (28) computes cell-type fractions optimized for tumor microenvironment characterization, and xCell (29) provides gene-signature enrichment scores for 64 cell types. Output includes per-sample stacked bar charts, per-cell-type box plots, and group-level Wilcoxon rank-sum comparisons with Benjamini-Hochberg correction.

## Regulatory analysis

### Hybrid gene regulatory network inference

Gene regulatory networks model the directed regulatory relationships between transcription factors and their target genes, providing a mechanistic context for expression changes and pointing to potential therapeutic targets through master regulator analysis (8). TransXplorer infers these relationships through a hybrid approach that balances confidence with organism coverage.

DoRothEA (11) curates TF-target interactions from gene-expression signatures, motif analysis, ChIP-seq binding, and literature, assigning each a confidence level (A: ChIP-seq-validated direct binding; B: ≥2 independent evidence types; C: single computational method; D: lower confidence). TransXplorer uses levels A–C to balance coverage and reliability. DoRothEA covers human and mouse; TFLink (10) extends coverage to rat, zebrafish, Drosophila, C. elegans, S. cerevisiae, Arabidopsis, chicken, pig, and bovine via predicted associations derived from literature mining and computational inference.

Master regulator analysis ranks transcription factors by the number of differentially expressed target genes they regulate, identifying candidates whose modulation could produce broad downstream effects. For human and mouse data, the hybrid mode prioritizes DoRothEA interactions supplemented with TFLink predictions; for all other organisms, TFLink serves as the primary source.

DEG threshold convention. Functional enrichment, PPI hub detection, and drug-target mapping use a stringent threshold of |log2FC| > 2 with adjusted P < 0.05 to focus on biologically robust changes. For master-regulator analysis only, this is relaxed to |log2FC| > 1 (adjusted P < 0.05) to preserve sufficient per-TF target coverage, since viper’s normalized enrichment scores become unstable when regulons contain too few responsive targets, a recommendation consistent with the DoRothEA developers’ guidance for master-regulator inference. The two thresholds are applied within a single analysis run and are reported explicitly in the case-study text and figure legends.

## Translational analysis

### TCGA integration and clinical validation

The platform uses TCGAbiolinks (30) to access RNA-seq data from 33 TCGA (The Cancer Genome Atlas) cancer cohorts. This allows for the identification of prognostic associations for genes of interest through Cox proportional hazards regression, Kaplan-Meier survival analysis with log-rank testing, and tumor versus normal differential expression analysis. For gene-level boxplots and survival curves, pre-calculated expression matrices offer query response times of less than a second.

### Real-time drug-target discovery and prioritization

Converting differential expression results into therapeutic hypotheses normally requires researchers to search pharmacological databases individually, cross-reference with clinical evidence, and assess target druggability. TransXplorer automates this process by coordinating queries against two complementary resources.

The Drug Gene Interaction Database (DGIdb) (12) aggregates drug-target relationships from over 40 sources, including ChEMBL (31), DrugBank, PharmGKB, the Therapeutic Target Database, and CIViC. TransXplorer submits differentially expressed genes to this pipeline, which retrieves interaction types, drug names, approval statuses, and source-database provenance. For genes with DGIdb hits, the OpenTargets Platform (13) is cross-referenced to obtain clinical trial data, disease associations, and tractability assessments. Using both sources together filters for candidates supported by both mechanistic drug-target evidence and clinical validation.

Three analysis modes serve different research objectives: Drug Discovery (DGIdb/ChEMBL focus), Disease Context (OpenTargets clinical focus), and Comprehensive (cross-validated) mode. Results are ranked by a prioritization score incorporating clinical development phase, number of supporting sources, cross-database concordance, and evidence quality. Targets appearing in both DGIdb and OpenTargets are flagged as high-confidence candidates.

### Universal organism support

TransXplorer pre-installs annotations for 11 commonly studied organisms (human, mouse, rat, Drosophila, zebrafish, C. elegans, yeast, Arabidopsis, chicken, pig, cow). For organisms beyond this set, Bioconductor’s AnnotationHub (32) supplies OrgDb packages for >1,800 species, downloaded on demand for gene-ID conversion and functional enrichment. KEGG pathway analysis covers ∼8,000 organisms, and GO enrichment is supported for any organism with an OrgDb entry. Cell-type deconvolution (human only), drug-target mapping (human only), and DoRothEA-based regulation (human and mouse) are auto-disabled for unsupported organisms, with the interface indicating which analyses remain available.

### Technical architecture

The platform is implemented in R Shiny and installed on a dedicated server (AMD EPYC 7351P, 32 threads, 125 GB RAM) in a Docker container behind an Nginx reverse proxy. The UI remains responsive through lazy loading of analytical modules and asynchronous processing. WGCNA was finished in less than eight minutes, drug-target queries for 500 DEGs in less than ninety seconds, and differential expression on 45,000 genes across 124 samples in less than three minutes.

Sessions can be saved and resumed across browsers for up to seven days using run IDs.

## RESULTS

The platform’s analytical scope is demonstrated by two use cases. In the first, the entire pipeline, from quality control to drug-target mapping, is applied to a dataset on cardiac endothelial differentiation (GSE151427). The second examines TCGA kidney renal papillary cell carcinoma (KIRP) and uses survival and expression data to verify a candidate gene.

### Use Case 1: Cardiac endothelial differentiation (GSE151427)

#### Data loading and quality control

Dataset GSE151427 compares cardiac mesoderm-derived endothelial cells (CMEC) and paraxial mesoderm-derived endothelial cells (PMEC) from human induced pluripotent stem cells. It contains 22 samples from differentiation days 6 and 8. The timepoint variable introduces potential batch structure.

#### Automated batch effect detection and correction

“Time” was identified by the batch evaluation as a confounding variable (composite score 0.559; PVCA: 34.1%, kBET rejection rate: 0.473, Silhouette: 0.375), above the 0.25 corrective threshold (see Supplementary Figure S1). The Silhouette score was automatically corrected using Limma’s removeBatchEffect, resulting in a 58.7% decrease from 0.375 to 0.155. PCA classified data by differentiation time point before correction; following correction, CMEC and PMEC separated along PC1. This was verified by UMAP (Figure 1A). A researcher who was not aware of the confounding variable would have incorrectly analyzed the dataset without an automated detection tool.

**Figure 1.**
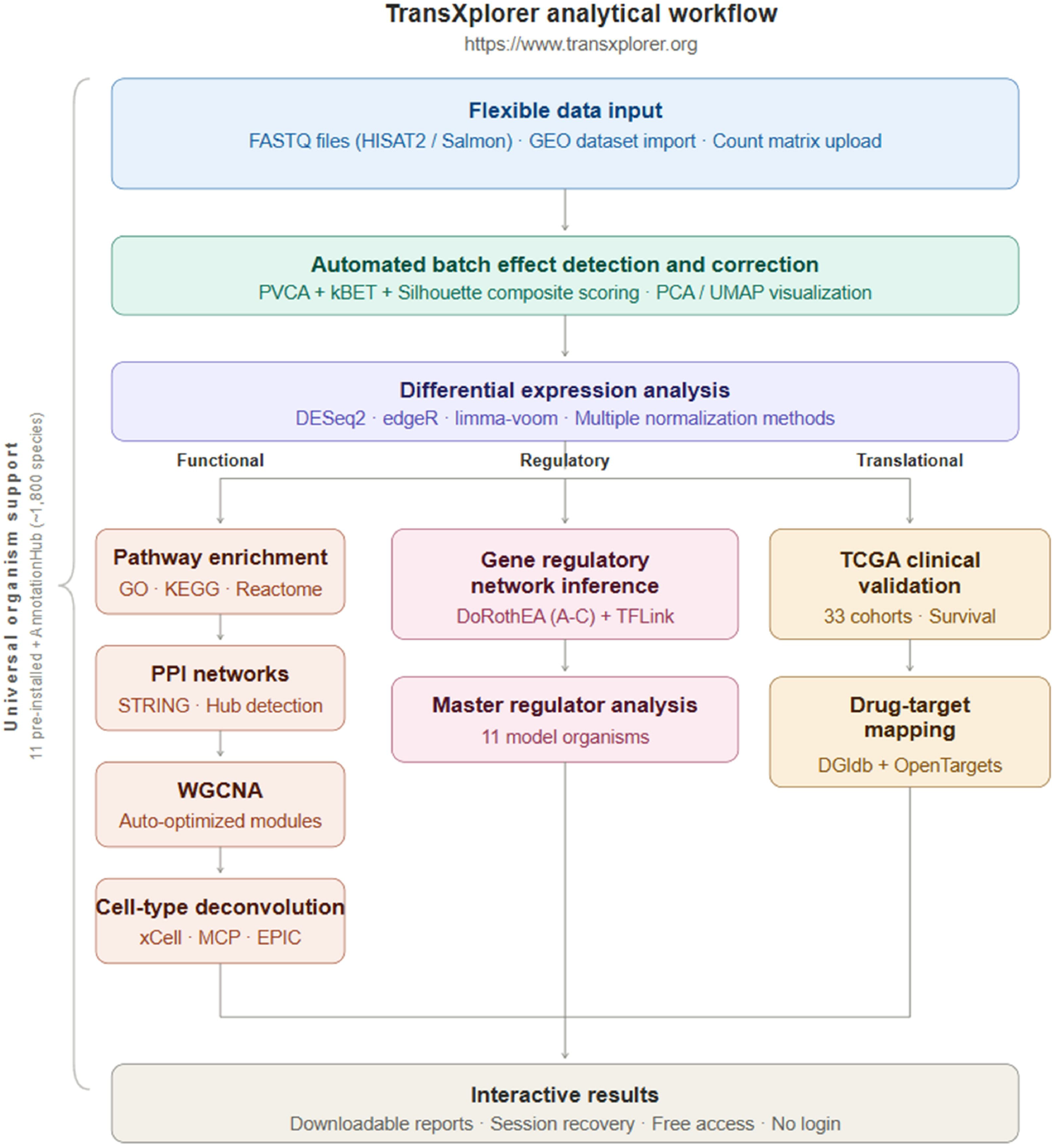
TransXplorer analytical workflow. Schematic of the end-to-end RNA-seq analysis pipeline available at https://www.transxplorer.org. Three flexible inputs feed the platform: paired/single-end FASTQ files (aligned with HISAT2 or pseudo-quantified with Salmon), direct GEO Series import (e.g., GSE164073 resolved automatically through GEOquery), or a pre-computed count matrix. All inputs flow through an automated quality-control stage that combines PVCA, kBET, and Silhouette scoring into a single composite metric; when the score exceeds 0.25, the platform corrects the largest confounding variable with limma’s removeBatchEffect and reports PCA / UMAP projections before and after. Differential expressions run in DESeq2, edgeR, or limma-voom with user-selected normalization (TMM, RLE, VST, CPM, or quantile). DEG output dispatches to three parallel analytical limbs: a Functional branch (clusterProfiler ORA + GSEA over GO/KEGG/Reactome → STRING-based PPI with hub detection → auto-optimized WGCNA → cell-type deconvolution via xCell, MCP-counter, and EPIC), a Regulatory branch (gene regulatory network inference using DoRothEA confidence levels A–C augmented with TFLink → master-regulator ranking and viper-based TF activity inference across 11 model organisms), and a Translational branch (clinical validation across all 33 TCGA cohorts with survival analysis → drug-target mapping that combines DGIdb and OpenTargets). Universal organism support (≈1,800 species via 11 pre-installed databases plus Bioconductor’s AnnotationHub on demand) is available throughout. All results are interactive and downloadable, and the sessions can be reopened from a unique run ID for up to seven days, after which the session is automatically deleted; no login is required.

#### Differential expression analysis

212 genes at |log2FC| ± 2 and adjusted p < 0.05 were found using edgeR with TMM normalization. CTHRC1, BOC, TNFRSF19, NR2F1, FZD10, and MAFB were among the upregulated genes; SMAD6, GATA4, BMPER, CRHBP, and GATA5 were among the downregulated genes (see Supplementary Figure S2A). These data can be viewed as a well-established volcano plot and heatmap for a simple visual approach, showing these important DEGs and the distinct separation of CMEC and PMEC in the top-20 DEG heatmap (see Supplementary Figure S2B).

#### Pathway enrichment analysis

**Over-representation analysis** via clusterProfiler, with all tested genes as background, returned 160 enriched pathways (adjusted p < 0.05) from GO, KEGG, and Reactome as selected databases for this run. Top terms were Heart Development (GO:0007507), Focal adhesion, ECM-receptor interaction, Regulation of Developmental Process (GO:0050793), Positive Regulation of miRNA Metabolic Process (GO:2000630), and Negative Regulation of Cell Differentiation (GO:0045596), matching the biology of two endothelial lineages at different specification stages (Figure 2C). Additionally, the pipeline creates an interactive pathway-gene network (Figure 2D) with 85 edges connecting 18 routes to 49 genes. This kind of visualization aids in cross-pathway database verification as well as identifying which DEGs are more involved in different pathways.

**Figure 2.**
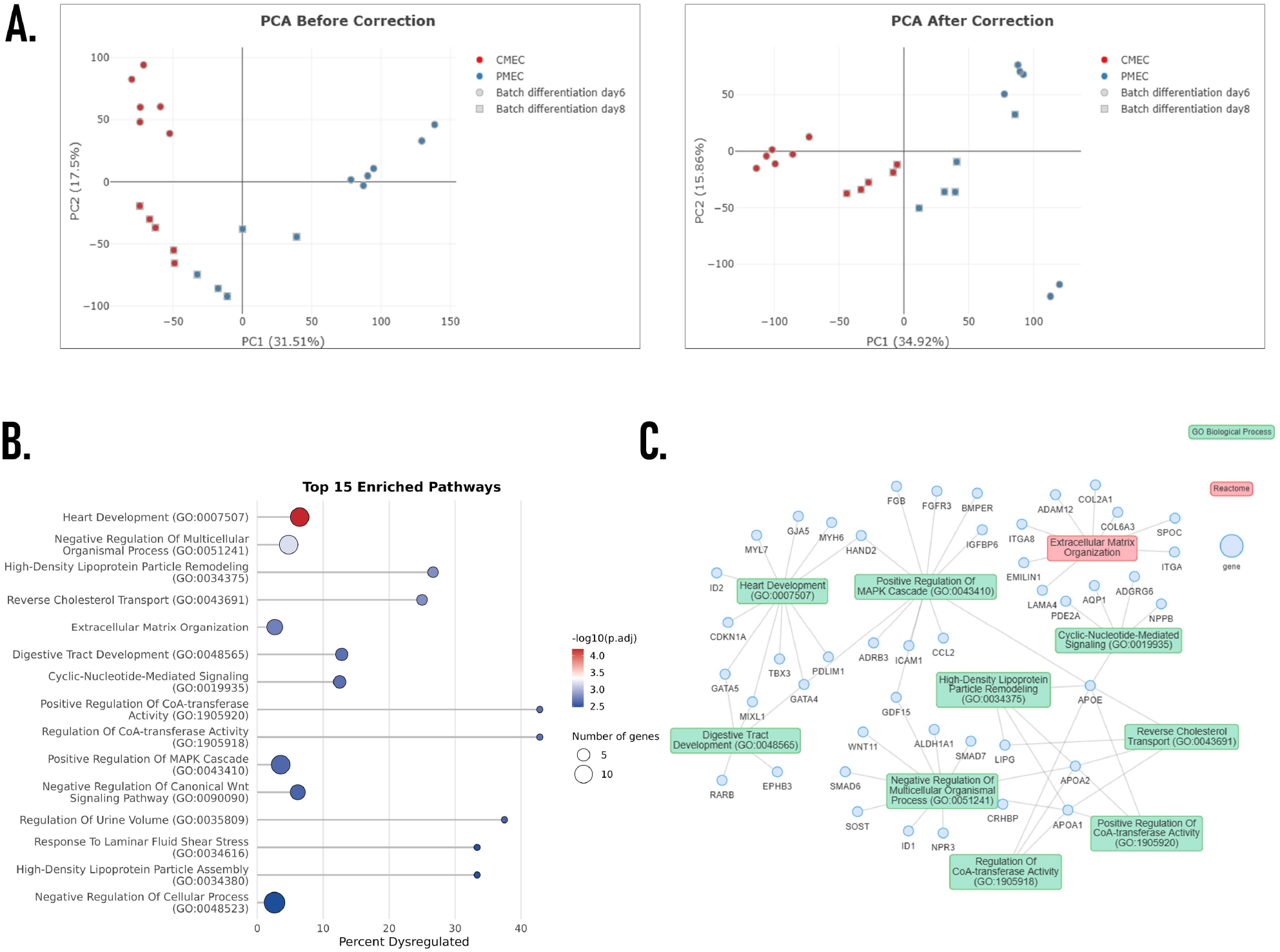
Functional analysis of cardiac vs paraxial-mesoderm-derived endothelial cells (GSE151427, n = 22). Demonstration of the Functional analytical branch on iPSC-derived cardiac mesoderm-derived endothelial cells (CMEC) versus paraxial mesoderm-derived endothelial cells (PMEC) sampled at differentiation days 6 and 8 (11 samples per condition). (A) PCA projections before (left) and after (right) automatic batch-effect correction. The composite QC score for the “Time” variable was 0.559 (PVCA 34.1 %, kBET rejection 0.473, Silhouette 0.375; threshold 0.25), triggering correction with limma’s removeBatchEffect. Pre-correction samples cluster by differentiation day (circles = day 6, squares = day 8); post-correction, they separate by biological condition along PC1 (CMEC red, PMEC blue), with Silhouette dropping 58.7 % (0.375 → 0.155). Two PMEC samples (blue circles, bottom-right of the post-correction panel) are day-6 collections that retain residual differentiation-stage separation along PC2 after batch correction; limma’s adjustment removes technical-batch covariance only, preserving genuine time-course biology. (B) Lollipop plot of the top 15 enriched GO Biological Process and Reactome pathways from clusterProfiler ORA on the 212 DEGs that pass |log_2_FC| > 2 and adjusted P < 0.05. The stem extends to the percentage of pathway genes recovered as DEGs (x-axis); bubble size encodes DEG count, and bubble colour encodes -log_10_(adjusted P) on a blue-white-red gradient. Heart Development (GO:0007507) is the most significant term, alongside HDL particle remodelling, Reverse Cholesterol Transport, and Extracellular Matrix Organization, all consistent with cardiac endothelial specification. (C) Bipartite pathway-gene network (cnetplot) connecting 18 enriched pathways (GO Biological Process: green; Reactome: red) to 49 DEGs (blue) through 85 edges. Visualization exposes cross-database overlap (e.g., GATA4 and HAND2 contributing to multiple cardiac-development terms) and identifies DEGs whose dysregulation has the broadest pathway footprint.

#### Cell-type deconvolution

MCP-counter estimated endothelial cells at **58.18%** (p = 5.17 × 10^−11^) and fibroblasts at **29.04%** (p = 2.87 × 10^−11^), consistent with the iPSC-to-endothelial differentiation protocol used in this dataset. CMEC samples scored higher for endothelial cells, while PMEC had elevated mesenchymal stromal and fibroblast proportions (see Supplementary Figure S3A–B). This difference is expected: paraxial mesoderm gives rise to a broader range of connective tissue lineages than cardiac mesoderm, and prior studies have reported a larger stromal fraction in paraxial mesoderm-derived cultures (33). The platform’s ability to recover this known lineage difference from bulk expression data without prior knowledge of the differentiation protocol supports the reliability of the deconvolution module.

#### Protein-protein interaction network

A connected network of 158 proteins connected by 884 interactions was obtained from STRING-based PPI analysis on the 211 differentially expressed input genes, covering 74.9% of the input set (see Supplementary Figure S4A for the network plot). The degree distribution was skewed to the right (see Supplementary Figure S4C), a topology typical of biological networks where large-scale functional programs are coordinated by a small number of highly linked nodes. CCND1 (cyclin D1, degree 52) was the top hub by degree, followed by APOE (46), ICAM1 (42), and GATA4 (42); Supplementary Figure S4B displays the whole top 15 ranking. Among these major hubs, GATA4 is one of the first and most researched cardiac transcription factors; it is necessary for cardiomyocyte specification during embryonic development and directly activates cardiac structural genes (MYH6, TNNT2, NPPA) (34). Its function in defining cardiac lineage identity is consistent with its emergence as a significant hub in a comparison between cardiac and paraxial-endothelial cells. Thirteen cardiac transcription factors and structural genes (GATA4, GATA5, TBX3, HAND2, MYH6, MYL7, GJA5, APLN, PDLIM7, FOXC1, ID2, MIXL1, CDKN1A) supported the network’s dominant functional enrichment (see Supplementary Figure S4D) (GO:0007507, adjusted p = 1.04 × 10^−9^).

#### Gene regulatory network analysis

Hybrid GRN inference using DoRothEA (levels A–C), applied to 778 DEGs (|log2FC| > 1, adjusted P < 0.05; relaxed from the |log2FC| > 2 threshold used for pathway enrichment, PPI, and drug-target analyses to ensure sufficient per-TF target coverage for viper-based master-regulator analysis), produced a directed regulatory network of 2,758 activator/repressor edges between 27 transcription factors and 369 of the input DEGs (47% DEG coverage), with 262 TF–TF regulatory links connecting the master regulators in a layered hierarchy (Figure 3A). 87% of edges (2,417 activating, 376 repressing) were classified as activating, consistent with a transcriptional-activation-dominated programme during early lineage specification. The top five master regulators ranked by composite influence score (target count, evidence confidence, and ChEMBL druggability) were MYC (386 targets; score 70.3), E2F1 (324; 65.3), EGR1 (124; 45.7), ETS2 (30; 37.9), and SMAD3 (33; 37.0); the broader 10-TF master-regulator set further included cardiac and vascular regulators TEAD4, NR2F1, NFATC1, MEIS1, and GATA6 (Figure 3B). The cell-cycle signature observed in the WGCNA brown and yellow modules and in the cyclin-anchored PPI hubs (CCND1, CDKN1A) is recapitulated by the MYC and E2F1 dominance, which is compatible with the actively dividing condition of iPSC-derived endothelial cells in culture. Several cardiac and vascular regulators co-emerged in the same top tier: MEIS1, which patterns the second heart field; NR2F1 (COUP-TF1), which specifies arterial–venous endothelial identity (35); TEAD4, the Hippo-pathway effector active in vascular development; NFATC1, which is necessary for endocardial cushion and heart-valve morphogenesis (36); and GATA6, a core cardiac transcription factor that, with GATA4, drives cardiomyocyte specification (37,38). The presence of multiple GATA-family regulators in this tier also tracks with the chromatin-level landscape of cardiac transcription-factor activity reported by Paige et al. (39). Their combined enrichment as master regulators in a comparison of PMEC and CMEC is biologically consistent with both lineages’ shared endothelial program and CMEC’s cardiac-mesoderm origin. Viper-calculated TF activity scores (Figure 3C) further resolve the separation: cardiac/lineage TFs (GATA2, NR2F1, TWIST1, EGR1, SMAD3) and cell-cycle regulators (MYC, MYCN, E2F1, TFDP1, LEF1) cluster in opposing directions across CMEC and PMEC samples, reproducing the lineage separation seen in PCA, deconvolution, and WGCNA module-trait analyses.

**Figure 3.**
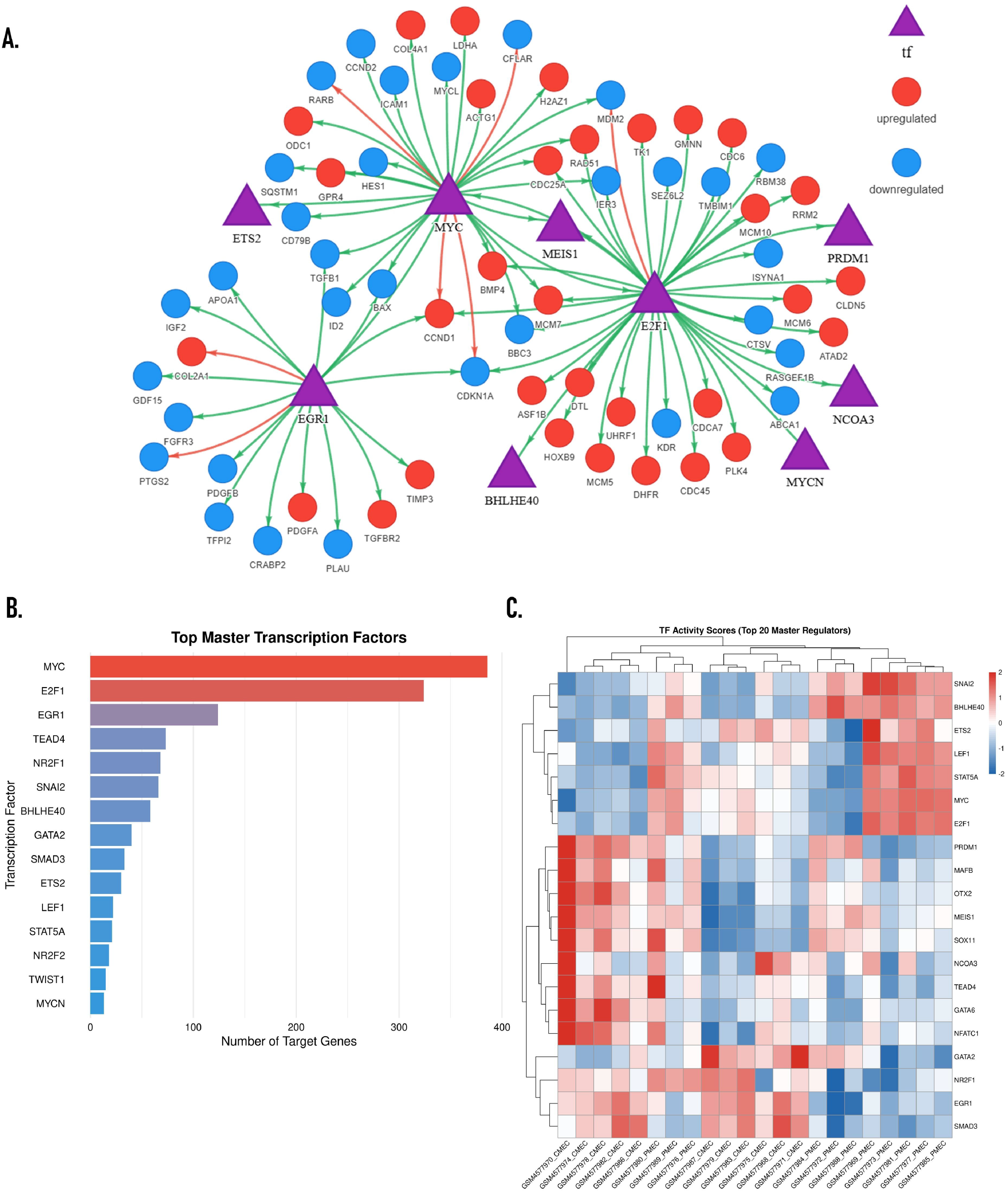
Regulatory analysis: Master transcription factor inference for CMEC vs PMEC (GSE151427). Demonstration of the Regulatory analytical branch. To preserve sufficient per-TF target coverage for stable viper-based inference, the GRN analysis uses a relaxed DEG threshold of |log_2_FC| > 1 and adjusted P < 0.05 (778 DEGs), versus |log_2_FC| > 2 used elsewhere. Hybrid GRN inference using DoRothEA confidence levels A-C produced a directed network of 2,758 regulatory edges (87% activating) between 27 transcription factors and 369 of the 778 input DEGs (47% coverage), with 262 TF-TF regulatory links. (A) Master-regulator subnetwork showing the top transcription factors (purple triangles) and their direct target DEGs (circles), colour-coded by direction of dysregulation in CMEC vs PMEC (red = upregulated, blue = downregulated). Arrows indicate inferred regulatory edges from DoRothEA, with edge colour encoding the database’s mode of regulation (green = activation, red = repression). The subnetwork is anchored on the top 3 master regulators (MYC, E2F1, EGR1) selected as focus TFs; their downstream targets include DEG circles and additional purple triangles for TFs that are themselves DEGs (e.g. ETS2, MEIS1, BHLHE40, MYCN, NCOA3, PRDM1). The remaining members of the 10-TF master-regulator set are summarised in panel B. The subnetwork highlights TFs central to mesodermal patterning, cell-cycle control, and endothelial fate (MYC, E2F1, EGR1, ETS2, SMAD3) whose target sets span both up- and down-regulated DEGs, consistent with their role as bidirectional master regulators of lineage commitment. (B) Top master regulators ranked by composite influence score (target count, evidence confidence, ChEMBL druggability): MYC (386 targets; score 70.3), E2F1 (324; 65.3), EGR1 (124; 45.7), ETS2 (30; 37.9), and SMAD3 (33; 37.0) lead the 10-TF master-regulator set. Bar colour encodes dominant regulatory mode (red = predominantly activating; purple = mixed; blue = predominantly repressing). (C) Hierarchically clustered heatmap of viper-computed TF activity scores (z-scored across samples) for the same top master regulators identified in panel B, displayed across all 22 samples (column annotation distinguishes CMEC and PMEC). Red/blue cells indicate above/below-mean activity. Several master regulators are differentially active between CMEC and PMEC, confirming that the lineage difference is driven by condition-specific regulatory rewiring rather than by isolated expression-level shifts.

#### WGCNA co-expression analysis

Automated soft-threshold selection produced a satisfactory approximation of biological scale-free topology with a scale-free R2 of 0.67 (see Supplementary Figure S5A) at soft power 8, chosen in accordance with Langfelder and Horvath’s sample-size guideline for n < 30 unsigned networks. Seven co-expression modules were identified from 4,998 genes using hierarchical gene clustering with dynamic tree cutting (see Supplementary Figure S5B): black (2,105), brown (1,360), purple (504), yellow (458), magenta (388), midnightblue (85), and lightcyan (83) (see Supplementary Figure S5C). GSE1, CCDC69, and ZMAT3, proteins linked to chromatin remodelling, mitotic spindle assembly, and p53-regulated growth control, anchored the biggest module (black). Yellow’s hubs (CCNA2, AURKB, TPX2, NUF2, KIF23) defined an M-phase signature, whereas Brown’s hubs (MCM6, MCM10, USP1, FEN1, CLSPN) defined a DNA-replication signature. THSD1 and EFNB2, the standard arterial-endothelial marker, served as the anchors for purple. PLN (phospholamban) is a well-characterized cardiac-specific regulator of sarcoplasmic-reticulum calcium cycling, and its development as a hub in cells produced from cardiac mesoderm is biologically anticipated. Magenta, anchored by PLN, TECRL, and MYOM1, recapitulated the cardiac-mesoderm signature. Eigengene analysis was used to resolve module-specific expression patterns across samples (see Supplementary Figure S5D). Cell Cycle and Proliferation (best p = 5.08 × 10^−34^ in yellow), DNA, RNA, and Protein Homeostasis (best p = 8.44 × 10^−50^, dominated by Eukaryotic Translation Elongation), Organismal Systems (black), Immune and Inflammatory Response (purple and midnightblue), and Signaling Pathways (lightcyan) were all assigned by evidence-based module classification (see Supplementary Figure S5E). According to Mai et al. (40), iPSC-derived endothelial cells in culture are actively proliferating, which is consistent with the prevailing cell-cycle and translation markers. The black (Pearson r = −0.94, p < 0.05) and magenta (r = −0.72, p < 0.05) eigengenes were strongly enriched in CMEC over PMEC, independently confirming the cardiac-mesoderm identity of magenta and the chromatin-remodelling program of black as CMEC-enriched; yellow (r = +0.92, p < 0.05) and purple (r = +0.84, p < 0.05) cell-cycle and vascular-maturation content. All seven modules exhibited functional enrichment.

#### Drug-target analysis

Comprehensive mode (DGIdb + OpenTargets) returned 402 compounds: 33 cross-validated in both databases, 334 with OpenTargets clinical evidence, 668 with OpenTargets interaction records (see Supplementary Figure S6A). The DGIdb table (see Supplementary Figure S6B) listed 534 entries with FGFR3 ranked highest (5 drugs, confidence 0.904), followed by TTR (3 drugs, 0.875) and RARB (5 drugs). The table provides information like drug name, interaction type, approval status, confidence score, evidence, and links to the original data sources for a detailed view of that drug.

OpenTargets, which provided information on Target druggability, clinical trials (Figure 4D), and disease associations, linked DEGs like ADRB3 to cardiovascular disease, KCNA5 to atrial fibrillation, and BTK to autoimmune conditions (see Supplementary Figure S6C).

**Figure 4.**
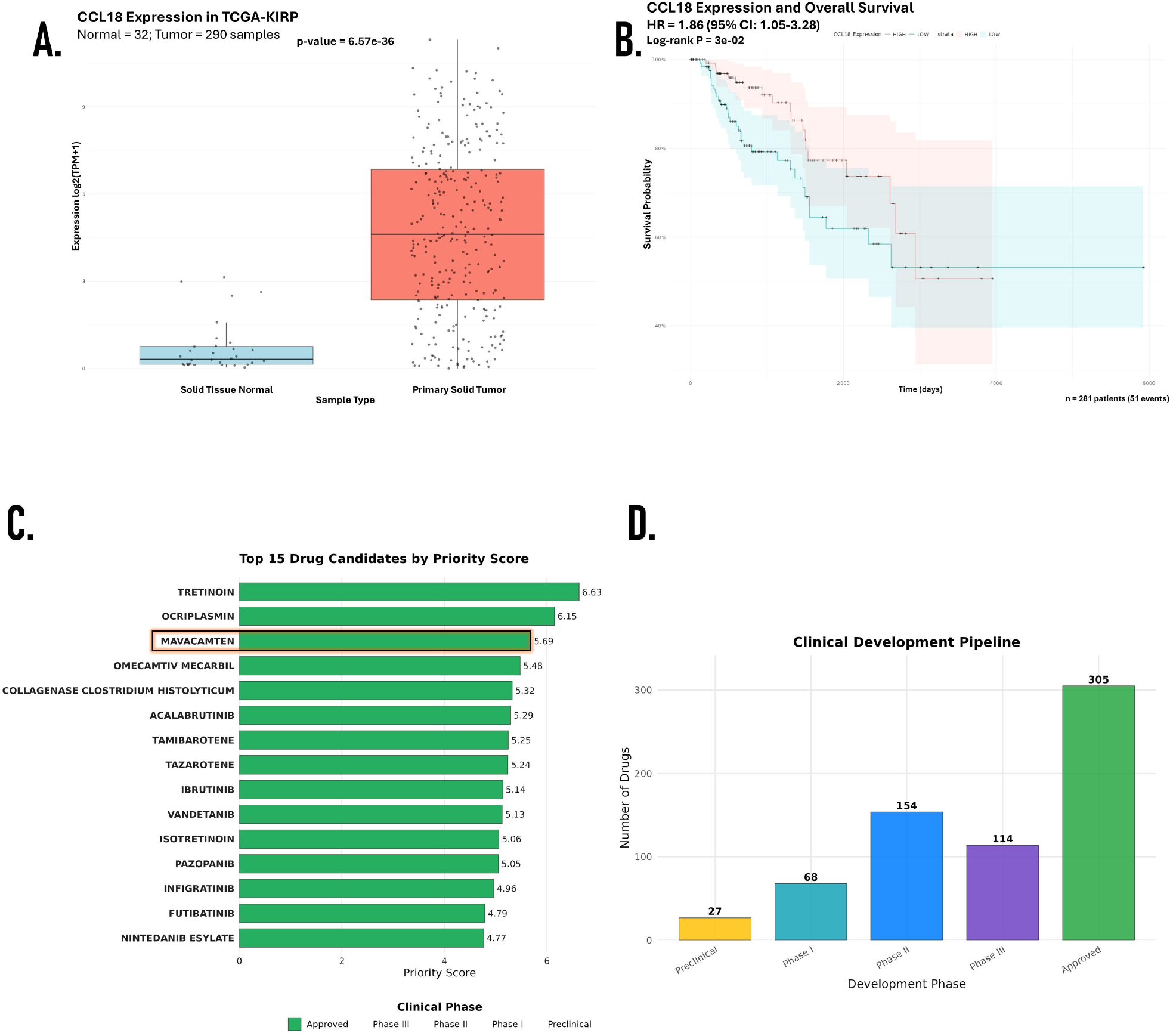
Translational outputs: Drug-target prioritization (CMEC vs PMEC) and clinical validation (TCGA-KIRP). Demonstration of the Translational analytical branch across both case studies. (A) Boxplot of CCL18 expression in TCGA-KIRP comparing primary tumour (n = 290) and adjacent normal tissue (n = 32); CCL18, surfaced as one of the most over-expressed DEGs from an unbiased differential expression analysis of 41,553 genes, is markedly elevated in tumour (Wilcoxon rank-sum P = 6.57 × 10^−36^; expression on log_2_(TPM + 1) scale). (B) Kaplan-Meier overall-survival analysis of the same KIRP cohort dichotomized at the median CCL18 expression. Patients in the high-expression stratum (n = 281, 51 events) show significantly worse outcomes than the low stratum (HR = 1.86, 95 % CI 1.05-3.28; log-rank P = 0.03), recapitulating CCL18’s published role as a tumour-associated-macrophage-derived prognostic marker and demonstrating that TransXplorer surfaces clinically relevant targets without prior hypothesis. (C) Top 15 drug candidates from the CMEC vs PMEC DEG list (|log_2_FC| > 2, adjusted P < 0.05; 212 DEGs) ranked by the composite TransXplorer Priority Score, which integrates bioactivity, clinical evidence, and target coverage from DGIdb and OpenTargets. Tretinoin scored highest (6.63; targets ALDH1A2, APOA1, MYL4, RARB), followed by Ocriplasmin (6.15) and Mavacamten (5.69; targets MYH6, MYL4, MYL7), the latter is an FDA-approved cardiac myosin inhibitor for hypertrophic cardiomyopathy, lending biological plausibility to the cardiac mesoderm-derived endothelial context. Bars are coloured by clinical phase (green = Approved). (D) Distribution of all 668 OpenTargets-mapped drugs for the CMEC vs PMEC DEG list across the clinical development pipeline (Preclinical 27 → Phase I 68 → Phase II 154 → Phase III 114 → Approved 305), showing that the platform surfaces both early-stage chemical probes and clinically deployed therapeutics for the same biology.

Lastly, in the drug ranking or drug prioritization table (Figure 4C), which was designed on multi-factor prioritization combining bioactivity, clinical evidence, and target coverage. In this table, we found drugs like Tretinoin placed first (score 6.63; targets ALDH1A2, APOA1, MYL4, RARB), then Ocrelizumab (6.15) and Mavacamten (5.89; targets MYH6, MYL4, MYL7). Mavacamten is a cardiac myosin inhibitor approved for hypertrophic cardiomyopathy, and its appearance here fits the cardiac endothelial context.

### Use Case 2: TCGA-KIRP RNAseq analysis and clinical validation

#### Dataset and differential expression

The TCGA-KIRP cohort (41,553 genes, 584 samples) yielded 2,447 DEGs by edgeR at |log2FC| >= 2 and adjusted p <= 0.05 (see Supplementary Figure S8A). The volcano plot (see Supplementary Figure S8B) and top 50 heatmap (see Supplementary Figure S8C) separate tumour, normal, and metastatic samples.

#### Pathway enrichment

Enriched pathways included immune cell infiltration, extracellular matrix remodelling, and metabolic reprogramming (see Supplementary Figure S8D). These match established features of the papillary RCC microenvironment: type 2 papillary tumours are characterized by dense immune infiltrates and a stroma-rich architecture driven by TGF-β signalling and collagen deposition (41). The identification of these pathways from an unbiased enrichment analysis, rather than from a curated gene set, supports the biological validity of the platform’s clusterProfiler implementation with proper background correction.

#### Gene-level clinical validation

One of the most elevated genes, CCL18, was compared to previously calculated TCGA expression data. Strong overexpression was seen in tumour tissue compared to normal (32 normal, 290 tumour; p = 6.57 × 10^−36^; Figure 4A). Additionally, a lower overall survival rate was linked to high CCL18 expression (HR = 1.86, 95% CI: 1.05–3.28, log-rank p = 0.03; 281 patients, 51 events; Figure 4B).

CCL18 is a chemokine secreted primarily by M2-polarized tumour-associated macrophages (TAMs). According to Chen et al. (42), CCL18 has been confirmed as an independent prognostic marker for breast cancer and promotes metastasis via the PITPNM3 receptor. Increased TAM density and a shorter disease-free survival are associated with higher CCL18 in clear cell RCC (43). Its emergence as the most elevated gene in KIRP, a kind of tumour where aggressive behaviour has been associated with macrophage infiltration, is biologically consistent and independently confirmed by published research. Notably, CCL18 was not selected from a candidate list; TransXplorer surfaced it as the highest-ranked gene from an unbiased differential expression analysis of 41,553 genes, demonstrating the platform’s capacity to identify clinically relevant targets without prior hypothesis.

### Comparison with existing platforms

Table 1 compares TransXplorer with four established RNA-seq web platforms. To the best of our knowledge, TransXplorer is the first platform that integrates FASTQ processing with dual quantification techniques, direct GEO dataset import, multi-organism regulatory network inference, real-time drug-target mapping, quantitative batch effect scoring with automatic correction, and universal organism support in a single environment. The largest pre-installed organism coverage (220 species) is provided by iDEP (4), which is devoid of regulatory networks, therapeutic mapping, and raw read processing. While NetworkAnalyst (9) offers network analysis, pharmacological mapping and differential expression require distinct techniques. Although GEPIA2 (5) does not process user-uploaded expression data or raw readings; it does facilitate TCGA-focused analysis. While interactive differential expression visualization is possible with DEBrowser (7), Network-level and translational analysis are not included.

**Table 1.**
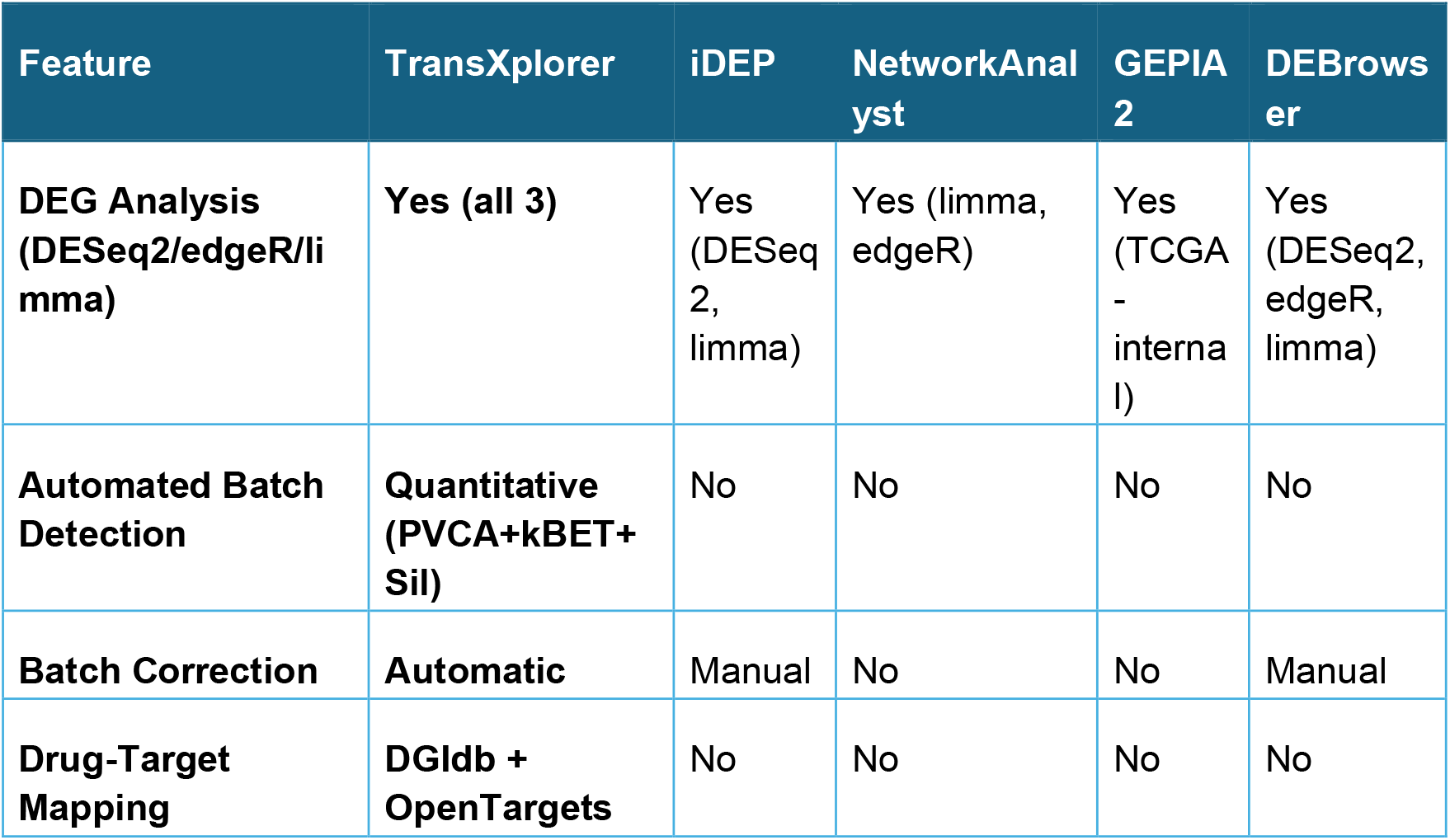
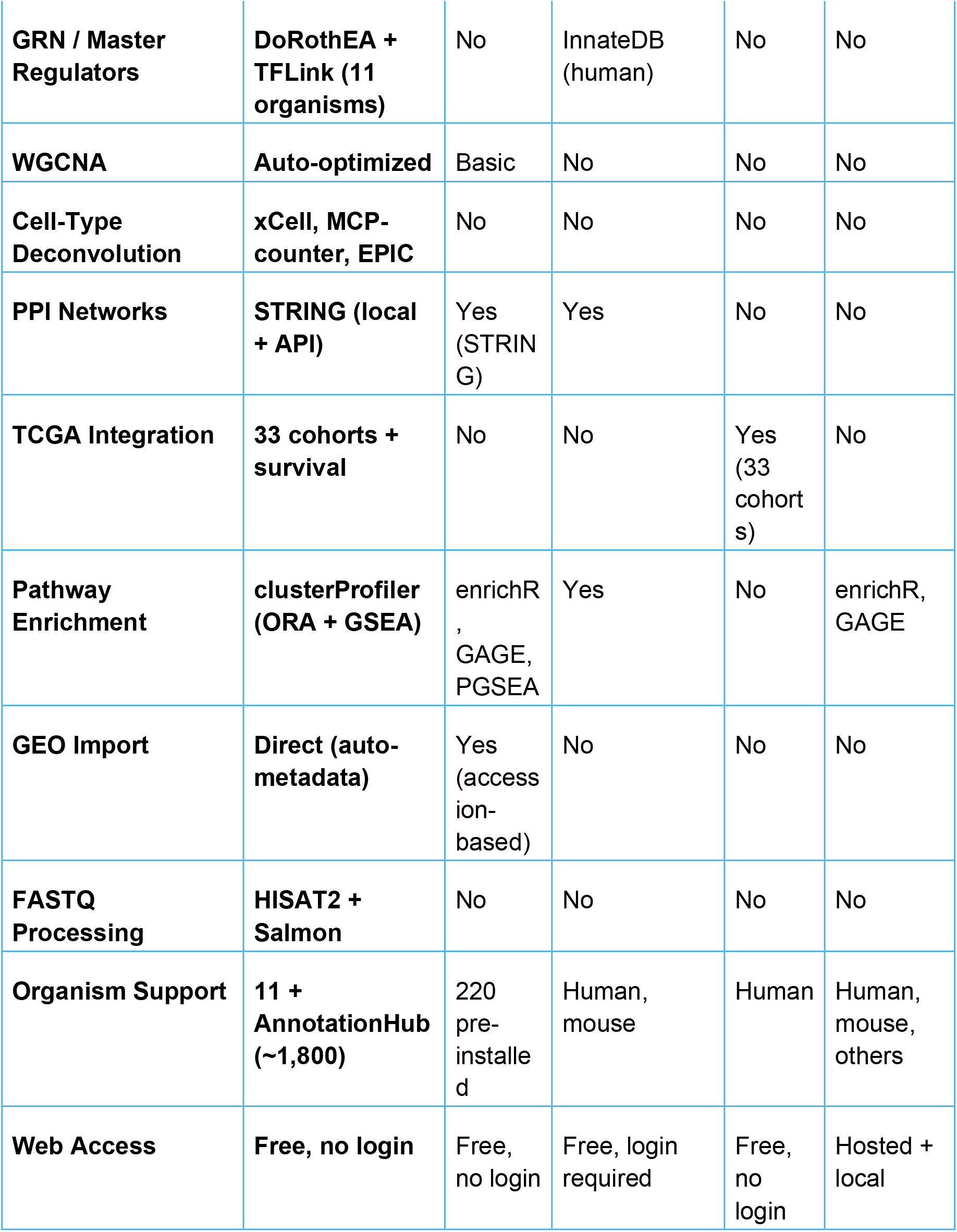
Feature comparison of RNA-seq web platforms.

## CONCLUSION

TransXplorer focuses on three practical issues in translational RNA-seq research: real-time translation of differential expression results into prioritized therapeutic candidates through integrated pharmacological database queries; automated batch confounding detection through quantitative multi-metric scoring; and regulatory network inference across diverse model organisms through hybrid curated-predicted transcription factor databases. For researchers who wish to work from raw readings or public data, the addition of direct GEO dataset import and an additional FASTQ processing pipeline with dual quantification methods (HISAT2 and Salmon) lowers the barrier.

TransXplorer, as seen from use case 1, is not only limited to the above-mentioned features but also provides daily-use, statistically and biologically more useful methods to get the most out of your data. Together with differential expression, pathway enrichment, co-expression networks, PPI analysis, cell-type deconvolution, and TCGA clinical validation, these features provide researchers without bioinformatics training a workable translational pipeline from raw data to therapeutic hypotheses. The platform is freely available at https://www.transxplorer.org with no login requirement.

## DATA AVAILABILITY

TransXplorer is freely available at https://www.transxplorer.org. No login is required. An example dataset (GSE151427) and tutorial documentation are provided within the application. The server has been tested on Chrome, Firefox, and Edge (Windows); Chrome (Linux); and Safari, Firefox, and Chrome (macOS).

The source code for the TransXplorer computational engine is available at https://github.com/varinder-madhav/transxplorer/ under the MIT License.

## ACKNOWLEDGEMENTS

We thank The Metabolomics Innovation Centre (TMIC) Wishart node for providing server infrastructure to host the TransXplorer web platform at https://transxplorer.org. We are also grateful to members of the Mason lab for early testing and user feedback during development.

## SUPPLEMENTARY FIGURE LEGENDS

**Supplementary Figure S1. Batch-effect detection and correction (GSE151427)**. Composite quality-control score combining principal variance component analysis (PVCA), k-nearest-neighbour batch-effect testing (kBET), and Silhouette analysis identifies “Time” (differentiation day 6 vs 8) as the dominant confounding variable (composite 0.559; PVCA 34.1%, kBET rejection rate 0.473, Silhouette 0.375), exceeding the 0.25 corrective threshold. Limma removeBatchEffect drops the Silhouette score 58.7% (from 0.375 to 0.155); PCA/UMAP projections after correction separate samples by biological condition (CMEC vs PMEC) along PC1 rather than by collection time.

**Supplementary Figure S2. Differential expression analysis (CMEC vs PMEC)**. (A) Volcano plot of all 212 DEGs identified by edgeR with TMM normalization at |log2FC| > 2 and adjusted P < 0.05 (red = upregulated, blue = downregulated, grey = not significant). Labelled DEGs include CTHRC1, BOC, TNFRSF19, NR2F1, FZD10, and MAFB (up) and SMAD6, GATA4, BMPER, CRHBP, and GATA5 (down). (B) Hierarchically clustered heatmap of the top 20 DEGs across the 22-sample cohort; CMEC and PMEC form two clearly separated sample blocks.

**Supplementary Figure S3. Cell-type deconvolution**. Stacked-bar profiles of estimated cell-type fractions in (A) CMEC and (B) PMEC samples, derived from xCell, MCP-counter, and EPIC. CMEC samples score higher for endothelial cells, while PMEC samples show elevated mesenchymal stromal and fibroblast proportions, consistent with the broader connective-tissue lineage potential of paraxial mesoderm and reproducing a known biological difference (Orlova et al., 2014) without prior knowledge of the differentiation protocol.

**Supplementary Figure S4. STRING-based protein–protein interaction (PPI) network analysis**. (A) Connected component of the PPI network (158 proteins, 884 interactions) built from the 211 DEGs (74.9% coverage); node colour encodes log2FC and node size scales with degree centrality. (B) The top 15 hub proteins ranked by degree, led by CCND1 (degree 52), APOE (46), ICAM1 (42), and GATA4 (42). (C) Right-skewed node-degree distribution, typical of biological networks where a small number of highly connected hubs coordinate large-scale functional programs. (D) Top enriched GO Biological Process terms from clusterProfiler ORA on the network nodes; Heart Development (GO:0007507; adjusted P = 1.04 × 10^−9^) is the most significant term, supported by 13 cardiac transcription factors and structural genes (GATA4, GATA5, TBX3, HAND2, MYH6, MYL7, GJA5, APLN, PDLIM7, FOXC1, ID2, MIXL1, CDKN1A).

**Supplementary Figure S5. WGCNA co-expression analysis**. (A) Soft-threshold selection diagnostic showing scale-free topology fit (R^2^) and mean connectivity as a function of soft power; power 8 (R^2^ = 0.67) was selected following Langfelder and Horvath’s sample-size guideline for n < 30 unsigned networks. (B) Hierarchical gene-clustering dendrogram with dynamic tree cutting yielding seven co-expression modules, colour-banded at the bottom. (C) Module size distribution: black (2,105 genes), brown (1,360), purple (504), yellow (458), magenta (388), midnightblue (85), and lightcyan (83). (D) Module eigengene profiles across the 22 samples, separating CMEC- and PMEC-enriched modules. (E) Evidence-based module classification heatmap assigning each module to functional categories: Cell Cycle and Proliferation (yellow, brown), DNA/RNA/Protein Homeostasis (yellow), Organismal Systems (black, magenta), Immune and Inflammatory Response (purple, midnightblue), and Signaling Pathways (lightcyan).

**Supplementary Figure S6. Drug-target prioritization (CMEC vs PMEC)**. (A) Heatmap summarising comprehensive-mode results from DGIdb + OpenTargets on the 211 input DEGs: 402 unique compounds, of which 33 are cross-validated in both databases, 334 carry OpenTargets clinical evidence, and 668 have OpenTargets interaction records. (B) DGIdb interaction table (534 entries) with FGFR3 ranked highest (5 drugs, confidence 0.904), followed by TTR (3 drugs, 0.875) and RARB (5 drugs); columns include drug name, interaction type, approval status, confidence score, evidence, and source links. (C) OpenTargets indication categories linking DEGs to disease classes (e.g., ADRB3 → cardiovascular disease, KCNA5 → atrial fibrillation, BTK → autoimmune conditions).

**Supplementary Figure S7. Hybrid gene regulatory network supplement**. Extended view of the DoRothEA + TFLink master-regulator analysis used in Figure 3. The summary panel reports the 27-TF, 2,758-edge network with 47% DEG coverage (369/778 DEGs) and 262 TF-TF regulatory links forming a layered hierarchy; activating interactions account for 87% of edges (2,417 activating, 376 repressing). The TF regulatory hierarchy table lists the upstream regulators of each master TF (e.g., AR → MYC, AR → SNAI2, ARNTL → BHLHE40, CEBPA → MYC, CREB1 → EGR1), exposing the upstream nodes whose perturbation would cascade through the regulatory program identified in Figure 3.

**Supplementary Figure S8. TCGA-KIRP differential expression and clinical context**. edgeR-based differential expression on the TCGA-KIRP cohort (584 samples; 41,553 genes; |log2FC| ≥ 2 and adjusted P ≤ 0.05; 2,447 DEGs). (A) Sample-level overview/clustering separating tumour, adjacent-normal, and metastatic samples. (B) Volcano plot of all genes tested; CCL18 and other TAM/RCC-associated genes are among the most overexpressed. (C) Hierarchically clustered heatmap of the top 50 DEGs across all samples, showing tumour vs normal vs metastatic separation. (D) Top enriched pathways from clusterProfiler ORA: immune cell infiltration, extracellular matrix remodelling, and metabolic reprogramming, consistent with established features of the type-2 papillary RCC microenvironment.

